# Adolescent Cognitive Control, Theta Oscillations, and Social Motivation

**DOI:** 10.1101/366831

**Authors:** George A. Buzzell, Tyson V. Barker, Sonya V. Troller-Renfree, Edward M. Bernat, Maureen E. Bowers, Santiago Morales, Lindsay C. Bowman, Heather A. Henderson, Daniel S. Pine, Nathan A. Fox

**Affiliations:** University of Maryland, College Park, MD 20742; University of Oregon, Eugene, OR 97403; University of California, Davis 95616; University of Waterloo, Waterloo, ON N2L 3G1, Canada; Emotion and Development Branch, Intramural Research Program, National Institute of Mental Health (NIMH), Bethesda, MD 20814

**Keywords:** Theta, Cognitive Control, Adolescence, Social Motivation, Medial-Frontal Cortex (MFC), EEG, Time-Frequency, Error Monitoring, Proactive Control, Reactive Control

## Abstract

Theta oscillations (4-8 Hz) provide an organizing principle of cognitive control, allowing goal-directed behavior that is conserved across species. In human adults, theta power over medial-frontal cortex (MFC) underlies monitoring, whereas theta synchrony between MFC and lateral-frontal regions reflects control recruitment. Prior work has not separated theta before/after motor responses, nor explained how medial-lateral synchrony drives different kinds of control behaviors. Theta’s role during adolescence, a developmental window characterized by a motivation-control mismatch also remains unclear, preventing possible cross-species work. Here, adolescents performed a flanker task alone or under observation to increase social motivation. We separated theta dynamics immediately before/after motor responses, identifying functional dissociations. We also dissociate MFC connectivity with rostral/caudal frontal cortex and distinct forms of behavioral control, which further differed before/after response. Finally, social motivation was found to exclusively upregulate post-response error monitoring and changes in control to prevent future errors, as opposed to pre-response theta dynamics.

## Introduction

Goal-directed behavior in humans engages neurocognitive processes— commonly referred to as cognitive control—to coordinate system-level brain activity (Gratton, 2018). Cognitive control involves two primary components, including: 1) *monitoring* for conflict or errors, which indicate control is needed; 2) *control recruitment*, further broken down into *proactive control*, recruited before needed, and *reactive control*, recruited in a just-in-time manner (Braver, 2012). In various mammalian species, theta oscillations reflect organizing activity within a cross-level cognitive control system (Cavanagh and Frank, 2014; Cohen, 2017; Verguts, 2017) and time-frequency EEG analyses can map oscillations among brain regions in ways that are missed by other approaches, linking particular cognitive control subprocesses to neural oscillations. For example, in adults, increased theta power over medial-frontal cortex (MFC) underlies monitoring (Ullsperger et al., 2014), whereas theta connectivity between MFC and lateral-frontal regions reflects control recruitment (Cavanagh and Frank, 2014). Application of these methods to adolescent data provides a unique opportunity to inform mechanistic understandings of cognitive control during a critical period of development. Cognitive control develops throughout childhood to approach maturity in adolescence (Casey et al., 2001; Chatham et al., 2009; Luna et al., 2004), with motivational processes becoming uniquely salient during this period (Casey et al., 2008; Luciana and Collins, 2012; Nelson et al., 2005; Steinberg et al., 2008). Here, we leverage EEG and advanced time-frequency approaches to explore the role of theta oscillations and social motivation on the deployment of cognitive control systems during this vital window of human brain development.

Studying interactions between motivational and cognitive control processes could provide a more fundamental understanding of cognitive control (Botvinick and Braver, 2015; Cools, 2016; Holroyd and Yeung, 2012). While links between motivation and cognitive control have been well-studied at the *behavioral* level, the *neuroscience* of motivation-control interactions is quite limited (Botvinick and Braver, 2015), particularly in relation to social motivation. Social processes, such as observation and evaluation, can increase motivation (Chib et al., 2018; Crone, 2014), particularly during the adolescent period (Blakemore, 2008; Nelson et al., 2005). Recent work examines these issues (Barker et al., 2018; Buzzell et al., 2017a; Crone, 2014) with a limited range of neural techniques (fMRI, ERPs). However, few studies dissect the ways that social motivation influences particular cognitive control processes, particularly in relation to theta dynamics, which could serve as a bridge for cross-species work. Work in adults suggests that monetary incentives specifically affect *proactive* control (Botvinick and Braver, 2015), but comparable work is needed in adolescents focusing specifically on social motivation. Leveraging time-frequency approaches to parse cognitive control into particular subprocess (see Figure S1) allows for testing which aspects of cognitive control are influenced by social motivation.

Regarding cognitive control subprocesses, work in adults suggests that MFC monitors the need for control (Ullsperger et al., 2014), as reflected in increased theta power during conflict (conflict monitoring; Cohen and Donner, 2013) or errors (error monitoring; Cavanagh et al., 2009). Moreover, MFC appears to recruit lateral-frontal cortex (LFC) to instantiate top-down control (Miller, 2000; Miller and Cohen, 2001), observable as MFC-LFC theta synchrony after errors or in response to conflict (Bolaños et al., 2013; Cavanagh et al., 2009; Cohen and Donner, 2013). These findings have not been extended to adolescents, nor have they been used to examine effects of social motivation. Moreover, existing time-frequency work targets overall levels of LFC engagement. Theory based on fMRI evidence (Badre, 2008; Koechlin, 2003) suggests that MFC theta synchrony within particular LFC sub-regions may predict particular forms of post-error behavior. For example, more *rostral* regions of LFC, including dorsolateral prefrontal cortex (DLPFC) might relate more closely to post-error changes in selective attention and accuracy rates, whereas more caudal regions of LFC, including primary and pre-motor regions might link more closely to post-error changes in response times (see figure S1). Additionally, whereas *increased* post-error MFC-LFC connectivity may reflect proactive control designed to prevent future errors, *reduced* connectivity immediately before an erroneous response may reflect a reactive control failure that generates the error (Banich, 2009). The current report explores these predictions. Specifically, we leverage advanced time-frequency analyses to: 1) explore theta oscillations during adolescence, 2) elucidate novel cognitive control mechanisms, and 3) characterize heterogeneous influences from social motivation on particular cognitive control subprocesses.

## Materials & Methods

### Participants

The current report focuses on 144 adolescents (M age = 13.12 years, SD = .58, 73 male) that were part of a larger longitudinal study focused on socio-emotional development. Children were originally selected at 4 months-of-age based on observations of their behavior in the laboratory (Fox et al., 2001). The 144 participants reported here reflect adolescents who returned to the laboratory at approximately 13 years-of-age to perform a flanker task and had valid behavioral data; a total of 132 participants had valid EEG. A series of statistical analyses were performed on subsets of this sample following the removal of condition-specific outliers; EEG plots include all 132 participants with EEG data. Analyses of flanker task ERP data have previously been reported for a largely overlapping subset of these participants (Buzzell et al., 2017a), however, this report reflects the first investigation of time-frequency dynamics in this sample. All procedures were approved by the University of Maryland—College Park institutional review board; all parents provided written informed consent, and children provided assent.

### Flanker Task

In a counterbalanced order, participants completed a modified flanker task (Eriksen and Eriksen, 1974) twice; once while alone (nonsocial condition) and once while believing they were being observed by peers (social condition). During the nonsocial condition, participants were provided with computer-generated feedback following each block. Prior to completing the flanker task within the social condition participants logged into an intranet chatroom where they created a screen name and uploaded their picture while waiting for two other age-matched peers to also log into the chatroom (see Figure 1B). For the social condition, participants were led to believe that their flanker task performance would be monitored via webcam by these peers and that real-time feedback would be provided after each block. In reality, the peers were fictitious, and the feedback was computer generated. During the nonsocial condition, participants were instructed that no one would observe their performance and that the block-level feedback was simply computer generated (see Figure 1C). Prior work has established the validity of this paradigm and its ability to modulate social motivation (Barker et al., 2018).

**Figure 1.**
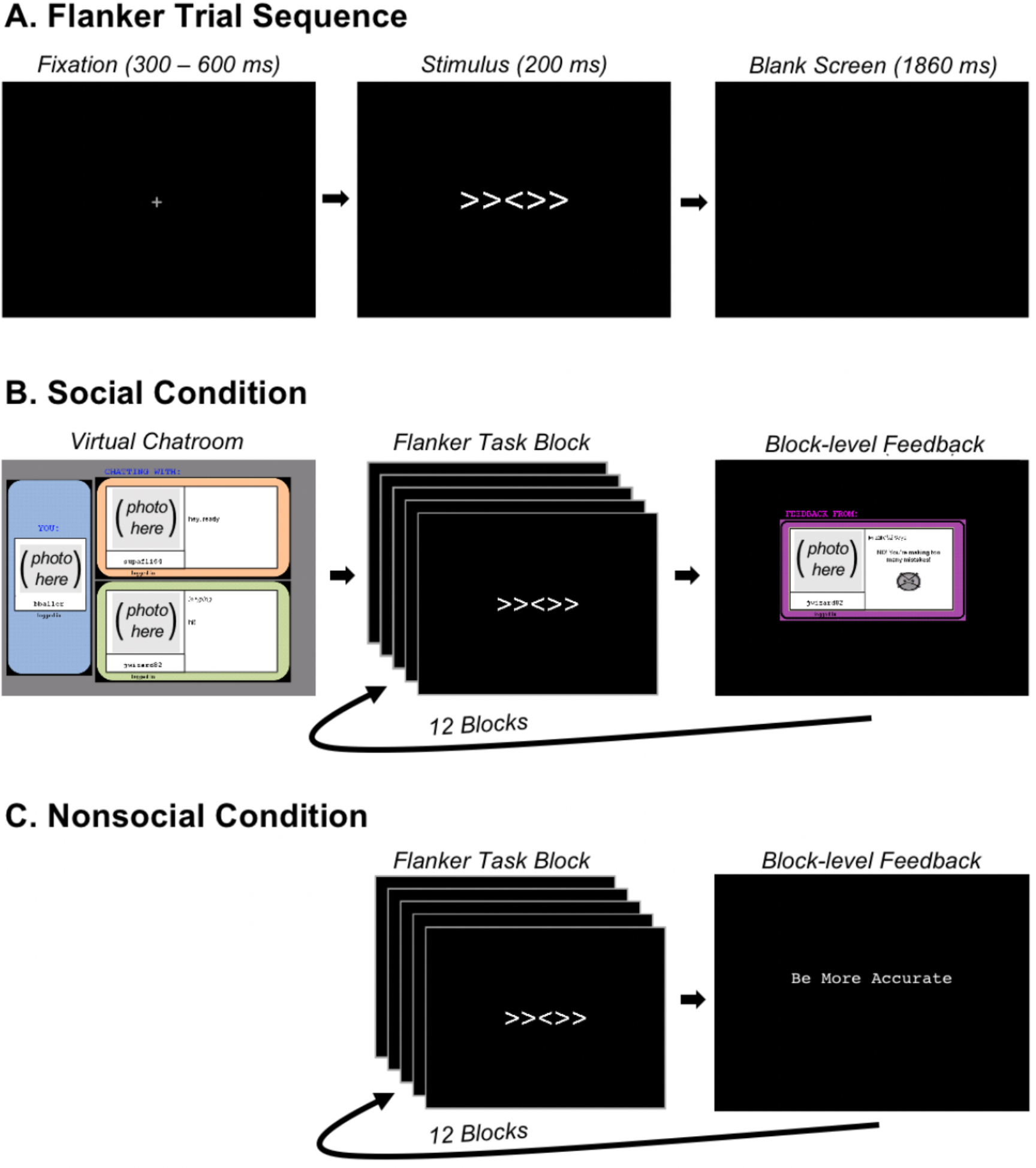
Experimental paradigm. A) Identical trial sequence employed within the social and nonsocial conditions (no trial-level feedback). B) Depiction of the virtual chat room and block-level social feedback employed within the social condition to increase social motivation. C) Depiction of the nonsocial condition in which block-level computer-based feedback was provided.

Within the social and the nonsocial conditions, block-level feedback always consisted of text indicating that the adolescent should “be more accurate”, “respond faster”, or that the adolescent was doing a “good job”. For the social condition, feedback consisted of both an emoticon and text (see Figure 1B), which the participant was led to believe was selected in real time by a peer. Nonsocial condition feedback consisted of text only. Although the adolescents believed that feedback was either provided by a peer or computer-generated, feedback was always generated based on task performance. If adolescents performed at or below 75% accuracy they received feedback indicating the need to “be more accurate”. If performance was at or above 90% they received feedback indicating the need to “respond faster”. If performance was between 75% and 90% they received feedback indicating that they were doing a “good job”. This feedback procedure is consistent with the recommendations by Gehring and colleagues (2012), which helped maintain accuracy at a level that would ensure an adequate number of errors for subsequent response-locked EEG analyses.

Each trial of the flanker task involved presentation of a central arrowhead flanked by two additional arrowheads on each side and facing in the same (congruent) or opposite (incongruent) direction (see Figure 1A). Participants were instructed to indicate the direction of the central arrowhead via button press and ignore the flanking arrowheads. Incongruent and congruent trials were presented with equal probability. For the social and non-social conditions, participants separately completed 12 blocks, each consisting of 32 trials and all blocks were followed by feedback. No feedback was presented at the trial level. Stimuli were presented on a 17” LCD monitor, using E-Prime 2.0.8.74 (Psychology Software Tools, Pittsburg, PA). Responses were collected using an EGI Response Pad (Model: 4608150-50) button box. The task was completed within a dimly lit, electrically-shielded and sound-attenuated room. Participants were left alone within the experimental room during data collection, with all monitoring of the task and EEG collection being performed within an adjacent room.

While prior work has validated that this task modulates social motivation (Barker et al., 2018), we additionally collected subjective reports of motivation from participants as a methods-check of our manipulation. Towards this end, participants were asked to rate on a scale of 1-10 “How hard did you try when you were playing the game that included feedback from the kids?”, as well as “How hard did you try when you were playing the game that included feedback from the computer?” Participants were further asked to provide a free-response explanation for their reported effort level in each condition. Statistical comparison of self-reported motivation between the social and non-social conditions was performed via a paired-samples t-test.

### EEG Acquisition and Preprocessing

EEG was acquired using a 128-channel HydroCel Geodesic Sensor Net and EGI software (Electrical Geodesic, Inc., Eugene, OR); EEG analysis was performed using the EEGLAB toolbox (Delorme and Makeig, 2004) and custom MATLAB scripts (The MathWorks, Natick, MA). Targeted electrode impedance level during data collection was < 50 kΩ, given that a high input-impedance system was used. Data were sampled online at 250 Hz and referenced to the vertex. Following acquisition, systematic marker offsets were measured and corrected for the EGI system (constant 36 ms offset) and E-Prime computer (constant 15 ms offset). Data were high-pass filtered at .3 Hz and low-pass filtered at 45 Hz. FAST tools (Nolan et al., 2010) were used to identify and remove bad channels. In order to identify and remove artifactual activity from the data, ICA decomposition was run on an identical data set with the addition of a 1 Hz high-pass filter (Viola et al., 2010). This 1 Hz filtered data set was epoched into arbitrary 1000 ms epochs; prior to running ICA, noisy epochs were detected and removed if amplitude was +/-1000 uV or if power within the 20-40hz band (after Fourier analysis) was greater than 30dB. If a channel led to more than 20% of the data being rejected, this channel was instead rejected. ICA was run on the 1 Hz high-pass filtered dataset and the ICA weights were then copied back to the original (continuous) .3 Hz high-pass filtered dataset (for an overview of this approach, see:Viola et al., 2010); all subsequent processing was performed on the .3 Hz high-pass filtered dataset. Artifactual ICA components were first detected in an automated procedure using the ADJUST toolbox (Mognon et al., 2011), followed by manual inspection of the ICA components. All ICA components identified as artifacts were subtracted from the data.

For EEG analyses, data were epoched to the response markers from -1000 to 2000 ms. All response-locked epochs were baseline corrected using the -400 to -200 ms period preceding the response. A final rejection of +/-100 uV was used to identify and remove bad epochs in the data that might have been missed by other methods. If greater than 20% of the data were rejected, the channel was rejected instead. All missing channels were then interpolated using a spherical spline interpolation. Following interpolation, data were referenced to the average of all electrodes. Given the focus on theta band activity, data were down-sampled to 32 Hz in order to improve computational speed with no loss to the signal of interest (i.e. theta = ~4-8 Hz; Nyquist = 16 Hz). All participants included in the EEG analyses had a minimum of 6 artifact-free trials, which has been shown to be suitable for response-locked analyses in either children or adults (Pontifex et al., 2010; Steele et al., 2016). However, to improve the robustness of the data and statistical analyses (Di Nocera and Ferlazzo, 2000), a bootstrapping approach was implemented. Specifically, for each participant, 4 trials were subsampled without replacement from the total number of trials 25 times to create an average EEG signal; these samples were bootstrapped 100 times to create the bootstrapped EEG signal that was utilized for all analyses and plotting. Note that bootstrapping also allows for matching trial numbers across conditions of interest.

### Isolation of Pre-and Post-Response Cognitive Control Within the Theta Band

#### Cohen’s class RID and time-frequency PCA

Using traditional time-frequency (TF) approaches, based on Morlet wavelets (Cohen, 2014; Herrmann et al., 2005), it is inherently difficult to separate pre-and post-response theta band activity. As a result, a common approach is to employ Morlet wavelets to yield a time-frequency decomposition, either for stimulus-or response-locked data, separately. Typically, only error-related effects are probed within response-locked data, only congruency-related effects within stimulus-locked data, and rarely are both error and congruency effects reported together for the same participants and task. However, this approach may miss unique effects manifesting immediately before versus after the response. Moreover, in studying stimulus-locked congruency effects, it is difficult to ascertain whether any theta dynamics truly reflect *pre*-response conflict monitoring and reactive control (to resolve conflict), as opposed to *post*-response error monitoring and proactive control prepare for the subsequent trial; a similar issue arises when attempting to interpret post-response theta and trying to rule out pre-response theta dynamics. To alleviate this issue, we first employed Cohen’s class reduced interference distributions (RID) to decompose a time-frequency representation of response-locked average power, which has proven superiority in its time-frequency resolution (Bernat et al., 2005); delta band activity was filtered out prior to TF decomposition to isolate theta activity. We then leveraged PCA of the average power time-frequency surface (TF-PCA; Bernat et al., 2005) to isolate separate sources of theta activity pre-and post-response. All conditions of interest (correct-congruent, correct-incongruent, error-congruent, error-incongruent) were entered into the PCA after first bootstrapping within conditions to match effect trial counts across conditions. See Table S1 as a reference for important terms and definitions.

Investigation of the scree plot suggested that a 3-factor solution described the data well. This 3-factor solution identified a clear MFC theta band factor maximal immediately following the response, consistent with prior work investigating post-error theta (Cavanagh et al., 2009), which has also been shown to contribute to the ERN (Luu et al., 2004; Trujillo and Allen, 2007). Critically, the 3-factor solution also yielded a pre-response theta factor with maximal activation over MFC and posterior scalp regions, consistent with prior work investigating stimulus locked theta for conflict (Cohen and Cavanagh, 2011; Cohen and Donner, 2013), which contributes to the N2 (Harper et al., 2014). A third alpha band factor was also identified, although investigation of this third factor is beyond the scope of the current report. See Figure 2 for a depiction of the pre-and post-response theta factors. These results represent the first evidence separating pre-and post-response MFC theta within the same epoch, removing confounds of pre-response theta on post-response theta, and vice-versa.

**Figure 2.**
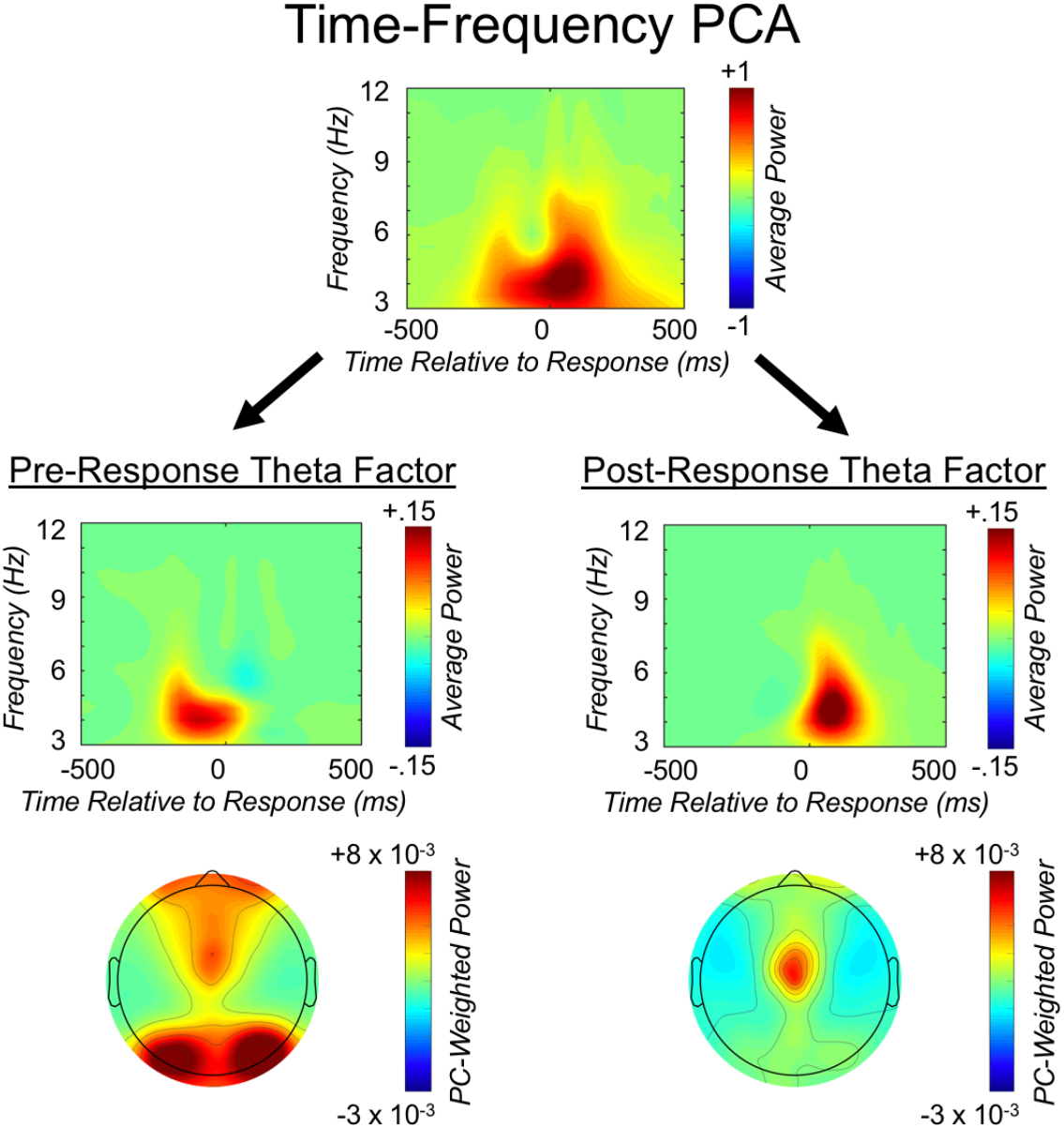
The Time-frequency PCA approach. We isolated separate pre- and post-response theta factors by applying time-frequency principle component analysis (PCA) to average power data. These factor loadings were subsequently applied to total power and also leveraged for inter-channel phase synchrony measurement. The top panel reflects the unweighted average power time-frequency distribution over medial-frontal cortex (MFC), collapsed across all conditions of interest. The second row depicts the same average power distribution weighted by the pre- and post-response theta factors, respectively; the third row displays the corresponding topographic plots.

#### Total power

After identifying pre- and post-response theta factors using the average power time-frequency surfaces, we then applied these factor loadings to a time-frequency decomposition of total power, again using Cohen’s Class RID and pre-filtering out delta. This approach allowed us to extract total power within the theta band immediately before, and immediately following the response; these data were used for subsequent analyses. Identifying the factor loadings first within the TF surface for average power improves separation of events occurring distinctly before and after the response; applying these loadings to a TF decomposition for total power incorporates both phase-locked and non-phase-locked data (Cohen, 2014), which is the most commonly employed metric for studying MFC monitoring processes and yields a more comprehensive measure of MFC theta (Cohen, 2014). Thus, we were able to extract a measure of total power that allowed for more direct comparisons with prior adult work, while still allowing us to retain the improved time-frequency resolution afforded by Cohen’s class RID and time-frequency PCA loadings (derived from the average power TF surface). In the text, all subsequent analyses and references to “theta power” refer to the total power measure weighted by the average power TF-PCA loadings. For analyses and plotting, MFC theta power was separately averaged for the pre- and post-response theta factors, within each condition of interest, for a cluster of electrodes that included E6 (approximately equal to FCz in the EGI geodesic sensor net) and the six immediately adjacent electrodes (E12, E5, E112, E106, E7, E13). Topographic plots exclude the outermost ring of electrodes, given that no a priori hypotheses related to these electrodes and they are subject to additional movement artifact; see Figure S2 for a complete map of electrode and cluster locations.

#### Inter-channel phase synchrony

Following the application of a Laplacian filter (current source density; CSD) to improve spatial resolution (Tenke and Kayser, 2012), we computed inter-channel phase-synchrony (across trials) to test for connectivity between an MFC seed electrode (FCz/E6) and six clusters over other cortical regions. Of note, raw EEG data is susceptible to volume conduction that degrades spatial resolution. However, a Laplacian transform mitigates volume conduction and enhances spatial resolution (Tenke and Kayser, 2012), allowing for the separation of the clusters tested here, and critically, distinguishing between rostral/caudal LFC. We applied the same pre- and post-response theta factor loadings to the connectivity surfaces in order to isolate pre- and post-response connectivity. We attempted to move beyond overall tests of MFC-LFC connectivity and leverage the improved spatial resolution afforded by a Laplacian transform. Specifically, we tested MFC connectivity with separate rostral and caudal LFC clusters, including: right (E123, E2, E122, E117, E124, E3) and left (E27, E26, E33, E28, E24, E23) rostro-LFC that most closely approximates DLPFC, as well as right (E103, E104, E110, E109) and left (E41, E36, E35, E46) caudal-LFC that most closely approximates primary and pre-motor cortex. We also included right (E90, E89, E83, E84) and left (65, 69, 70, 66) occipital regions in the connectivity analyses to test whether connectivity was indeed stronger for frontal areas relative to other brain regions. Topographic plots excluded the outermost ring of electrodes; see Figure S2 for a complete map of electrode and cluster locations.

### Experimental Design and Statistical Analyses

#### Overview

Prior research has not fully characterized the performance monitoring system in adolescence utilizing time-frequency analyses of EEG. Thus, we first sought to explore the neurobehavioral dynamics of the cognitive control system in the non-social condition. This supports further analyses probing social motivation (in the social condition) and provides results that can be more readily referenced to the broader performance monitoring literature in adults that typically lacks a social manipulation. Next, we computed difference scores for all behavioral and neural measures for the social and non-social conditions, separately, and ran an additional series of statistical analyses to probe the effect of social motivation on cognitive control. Collectively, this approach characterizes cognitive control dynamics in ways comparable to prior work, while facilitating analyses examining social motivation effects during this vital developmental period.

#### Statistical analyses

Data reduction and statistical analyses were performed using a combination of Matlab 2014b (The MathWorks, Natick, MA), R version 3.4.3 (R. Core Team, 2017), and SPSS version 24.0 (IBM Corp., Armonk, NY). Below, the complete details of each statistical analysis is described. For all analyses, alpha was set to .05; where appropriate, a Greenhouse-Geisser correction was employed for violations of sphericity, however, raw degrees of freedom are reported for ease of interpretation.

#### Behavioral data

For all behavioral analyses, response time (RT) analyses were restricted to correct trials and log-transformed prior to averaging. All analyses of RT data were performed on log-transformed data; raw RT values are reported in the text for ease of interpretation. Additionally, outliers (+/-3 SD) were removed from all conditions for both accuracy and RT measures; following the calculation of difference scores, outliers were again removed based on difference score values. We first calculated the overall accuracy and response times (RT) within the non-social condition. Next, we calculated the behavioral effects of stimulus conflict by calculating difference scores (incongruent minus congruent) for both accuracy and RT; for RT calculations, only correct trials were employed. These measures are referred to as “conflict-effect-accuracy”, and “conflict-effect-RT”, respectively.

In order to investigate behavioral changes following errors, we calculated accuracy rates and RT following error vs correct responses and difference scores were computed. In order to isolate post-error effects, congruency was held constant and only trials following error-incongruent and correct-incongruent trials were analyzed (with no congruency restriction for the post-error trial). We first calculated a behavioral metric indexing variation in task-specific proactive control, known as post-error reduction in interference (PERI; Danielmeier and Ullsperger, 2011). PERI was calculated by subtracting the conflict-effect-accuracy following correct responses from the conflict-effect-accuracy following errors. Thus, PERI indexes the degree to which task-specific control is instantiated following errors in order to *proactively* (Ridderinkhof et al., 2011) adjust selective attention (King et al., 2010) on the following trial, reflected by increased accuracy on the more difficult incongruent trials (relative to congruent). Next, we calculated the difference in RT on post-error vs post-correct trials, an effect known as “post-error slowing” (PES), thought to at least partially reflect a general and automatic inhibition of (pre-)motor cortex following unexpected events like errors (Danielmeier and Ullsperger, 2011; Jentzsch and Dudschig, 2009; Notebaert et al., 2009; Wessel, 2018; Wessel and Aron, 2017). Note that the deliberative and task-specific nature of PERI is in direct contrast to the more generalized and simple increase in RT (PES) following errors (Danielmeier and Ullsperger, 2011; King et al., 2010; Maier et al., 2011; Ridderinkhof, 2002). Below, we discuss testing for relations between error-related neural data and these post-error behavioral measures.

#### Error monitoring and post-error proactive control

We first tested whether MFC theta power increased following errors. Here, we held congruency constant (analyzing only incongruent trials) and employed a paired-samples t-test to compare error vs. correct differences within the post-response theta factor. Moreover, leveraging this same post-response theta factor and inter-channel phase-synchrony, we employed an ANOVA model to test whether a seed electrode over medial-frontal cortex (MFC) became more connected with electrode clusters over LFC following errors. For this model investigating post-error control recruitment, hemisphere (right, left), caudality (rostral-LFC, caudal-LFC, occipital) and accuracy (error-incongruent, correct-incongruent) were all entered as within-subjects factors. Collectively, this set of analyses tests whether theta oscillations relate to performance monitoring (increased theta power) and control recruitment (increased phase synchrony between MFC and LFC).

To confirm that MFC connectivity with LFC regions reflects the recruitment of transient *proactive* control for the *following* trial, we tested whether between-subject variation in MFC-LFC connectivity predicted between-subject changes in PERI. Here, our use of the term “proactive” control aligns with the dual mechanism of control (DMC) theory (Braver, 2012), suggesting control can either be recruited after the event requiring control in a “reactive” manner, or in preparation for an event requiring control in a “proactive” manner. Although DMC theory has most commonly been applied to reactive control at a transient, event-related level and proactive control at a sustained, block level, the proactive/reactive distinction also applies to transient changes in either proactive or reactive control (King et al., 2010; Ridderinkhof et al., 2011). PERI has been explicitly defined as a behavioral correlate of *transient proactive control following error commission* (Ridderinkhof et al., 2011) and linked to neural measures of selective attention (King et al., 2010). Within this view, PERI reflects an unequivocal and “ground-truth” measure of post-error *proactive* control. Thus, if increased MFC-LFC connectivity reflects transient proactive control increases (for the next trial), then we would expect individuals exhibiting the strongest increases in post-response MFC-LFC connectivity to exhibit the greatest increases in PERI. Moreover, based on theories specifying that more caudal-LFC regions are associated with relatively simple forms of control, whereas rostral-LFC regions are associated with more deliberative control (Badre, 2008; Koechlin, 2003), we hypothesized a similar dissociation would be observed here following errors. In particular, we predicted that MFC connectivity with *rostral*-LFC would predict PERI, a deliberative and task-specific form of proactive control. In contrast, we hypothesized that MFC connectivity with *caudal*-LFC regions would link to more automatic and general changes in behavior: PES. To test for these brain-behavior relations, post-response connectivity difference scores (error minus correct) were calculated for MFC connectivity with rostral- and caudal-frontal regions, outliers were removed, and we additionally collapsed across hemisphere prior to testing for relations with post-error behavior to reduce the number of comparisons; qualitatively similar relations were identified when not collapsing across hemisphere. Relations between these connectivity measures and both PES/PERI were tested using a family of Pearson’s product-moment correlation tests; a false-discovery rate (FDR) correction for multiple comparisons was applied using the method proposed by Benjamini and Hochberg (1995). We tested specificity in rostral vs. caudal connectivity predicting PERI by leveraging a Fisher’s r-to-z transformation and the method proposed by Steiger (1980); a similar comparison was performed for PES and rostral vs. caudal connectivity.

#### Conflict monitoring and pre-response reactive control

Within a speeded visuo-motor task like the flanker, errors can be prevented through two primary avenues: proactive or reactive control (Braver, 2012). As described above, error commission could drive a transient increase in proactive control to modulate behavior on the following trial (i.e. instantiated before the control is required). In contrast, reactive control is theorized to reflect the recruitment of control in a reactive and “just-in-time” manner as it is needed (i.e. *after* a stimulus is presented, but *before* the response). Thus, on a flanker task, one would expect a monitoring process to detect conflict and recruit reactive control following presentation of incongruent trials but prior to responding. Indeed, prior work (Cavanagh and Frank, 2014; Cohen and Cavanagh, 2011) has demonstrated that incongruent trials elicit an increase in monitoring (MFC theta power) and reactive control (MFC connectivity with LFC). However, we leverage the improved resolution of Cohen’s Class RID and time-frequency PCA to perform a separate test of pre-response monitoring and control, isolated from post-response cognitive control, all within the same epochs; this approach goes beyond prior analyses of peri-response theta in adults or adolescents. A conflict monitoring effect within the pre-response theta factor was tested via a paired-samples t-test comparing congruent-correct and incongruent-correct trials. An ANOVA model with hemisphere (right, left), caudality (rostral-LFC, caudal-LFC, occipital) and congruency (incongruent-correct, congruent-correct) as within-subjects factors investigated reactive control within the pre-response theta factor. Here, accuracy was held constant to isolate congruency effects. In line with a role in conflict monitoring, we hypothesized that MFC theta power would be increased for incongruent-correct (vs. congruent-correct) trials. Moreover, we hypothesized that incongruent-correct trials would be associated with increased pre-response connectivity between MFC and LFC, reflecting increased instantiation of reactive control in order to resolve the conflict associated with incongruent trials.

Additionally, we provide a third novel advance over prior work. If pre-response increases in MFC-LFC connectivity reflect the recruitment of reactive control in order to *prevent* an error, then *failures* of such control should be observable prior to error responses as a possible cause of the incorrect response. To test this claim, we investigated whether MFC-LFC connectivity within the pre-response theta factor was *reduced* prior to error responses using an ANOVA with hemisphere (right, left), caudality (rostral-LFC, caudal-LFC, occipital) and accuracy (error-incongruent, correct-incongruent) as within-subjects factors. Here, we held congruency constant, and analyzed only incongruent trials that are thought to require reactive control to resolve conflict and respond correctly. We hypothesized that pre-response MFC-LFC connectivity would be reduced prior to error-incongruent (vs. correct-incongruent) responses.

#### Cascade of pre- and post-response control

To summarize, our predictions for the series of connectivity analyses were as follows: 1) *Incongruent*-correct trials would yield a relative increase in pre-response MFC-LFC connectivity over *congruent*-correct trials, reflecting increased reactive control required to resolve conflict and respond correctly to incongruent stimuli; 2) Incongruent-*error* trials would exhibit reduced pre-response MFC-LFC connectivity compared to incongruent-*correct* trials, reflecting a failure of reactive control leading to an error response; 3) Following the response, *error*-incongruent trials would exhibit greater post-response increases in MFC-LFC connectivity compared to *correct*-incongruent trials, reflecting a transient increase in proactive control to adjust behavior for the following trial in order to prevent additional errors.

#### Testing effects of social motivation on cognitive control

Following analyses of the nonsocial condition characterizing theta band cognitive control dynamics during adolescence, we investigated how each of these processes were influenced by social motivation. Critically, identification of several distinct subprocesses underlying cognitive control allowed us to investigate possible heterogeneity in how social motivation influences particular subprocesses, as opposed to a monolithic influence. To reduce the number of comparisons and to avoid potential four-way interactions, we approached analyses of social motivation using difference scores for the behavioral and neural measures described above. Relevant difference scores were calculated based on accuracy or congruency for all behavioral and neural measures with outliers (+/-3 SD) being removed.

A series of paired-samples t-tests were employed to investigate behavioral differences as a function of social context. Similarly, separate paired-samples t-tests explored whether pre-response MFC theta power (conflict monitoring), or post-response MFC theta power (error monitoring) were influenced by social motivation. Finally, separate ANOVA models were employed to test whether pre- or post-response MFC-LFC connectivity was influenced by social motivation, as measures of reactive and proactive control, respectively. These ANOVA models were similar to those described above, with hemisphere (right, left) and caudality (rostral-LFC, caudal-LFC, occipital) as within-subjects factors, but also including social context (nonsocial, social) as a within-subject factor and employing connectivity difference scores as the dependent variable.

## Results

### Non-Social Condition

#### Behavior

For all analyses (behavioral and EEG), outliers (+/-3 SD) for specific conditions were removed where appropriate; each statistical analysis was performed using as much data as possible as opposed to listwise deletion across all analyses. Within the non-social condition, accuracy rates for congruent and incongruent trials were 95.11% and 74.47%, respectively; a one sample t-test for the conflict-effect-accuracy (incongruent minus congruent) was significant [*t*(1,143) = -26.91, *p* < .001]. Similarly, RTs for correct-congruent and correct-incongruent trials were 380.16 ms (SE = 3.77) and 454.22 ms (SE = 5.33), respectively; a one sample t-test for the conflict-effect-RT (incongruent minus congruent) was also significant [*t*(1,142) = 31.02, *p* < .001]. Collectively, this pattern of findings resembles results in prior research in both adolescents (Hogan et al., 2005) and adults (Botvinick et al., 2001). PES and PERI-accuracy were in the expected direction and above zero, although on average, there were no significant effects of PES [*t*(1,141) = 1.27, *p* = .207] or PERI-accuracy [*t*(1,142) = .9, *p* = .369] revealed by a pair of one-sample t-tests. Such null effects could arise from opposing individual differences in activation of post-response proactive control (MFC-LFC connectivity). That is, PES and PERI might only emerge for individuals exhibiting stronger MFC-LFC connectivity following errors; indeed, the following section reports significant relations between post-response MFC-LFC connectivity and these post-error behavioral metrics.

#### Error monitoring and post-error proactive control

A paired-samples t-test investigated post-response MFC theta power as a correlate of error monitoring; this test revealed an increase in total power for errors [*t*(1,127) = 11.48, *p* < .001]. The ANOVA investigating MFC connectivity within this same post-response theta factor revealed a main effect of accuracy [*F*(1,116) = 50.5, *p* < .001], caudality [*F*(2,232) = 99.45, *p* < .001] and a caudality-by-accuracy interaction [*F*(2,232) = 5.63, *p* = .004]. Follow-up comparisons, collapsing across hemisphere, demonstrated significantly stronger post-error connectivity between MFC and rostral-LFC regions, in comparison to occipital [*t*(1,116) = 3.29, *p* = .001]. MFC connectivity with caudal-LFC was also significantly stronger than occipital [*t*(1,116) = 2.39, *p* = .018], whereas connectivity magnitude did not differ between rostral- and caudal-LFC regions [*t*(1,116) = .62, *p* = .539]. This pattern of results is consistent with MFC-related monitoring (theta power) recruiting LFC-related control (MFC-LFC connectivity) following error commission. Figure 3 depicts the error/correct post-response theta power and connectivity results.

**Figure 3.**
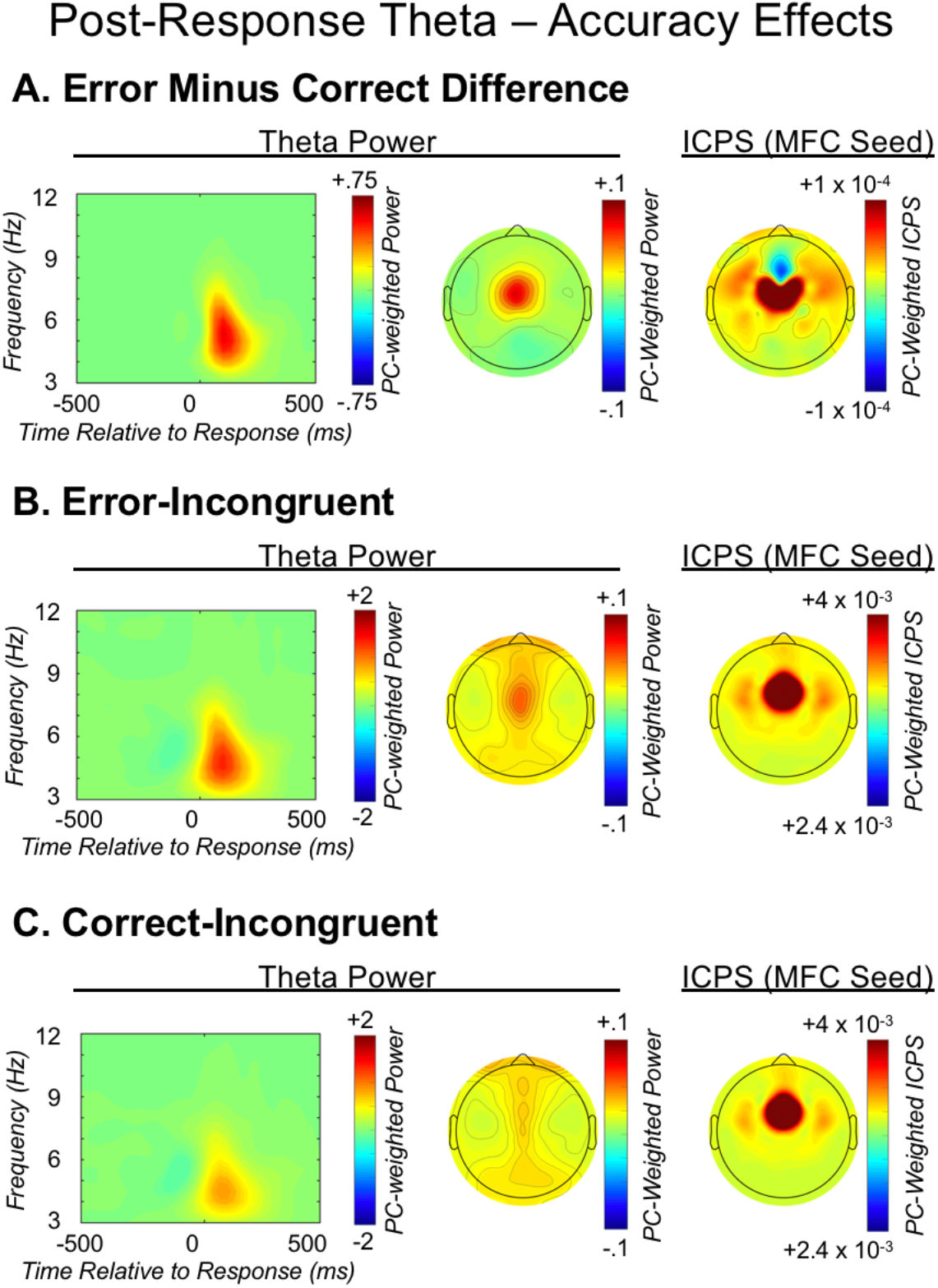
Post-response theta dynamics. From left to right, each row depicts: the medial-frontal cortex (MFC) total power time-frequency distribution weighted by the post-response theta factor; the corresponding topographic plot; MFC-seeded inter-channel phase synchrony (ICPS) within the post-response theta factor. The three rows present: A) the difference between error-incongruent and correct-incongruent activity; B) error-incongruent activity; C) correct-incongruent activity.

Task-specific improvement in accuracy following errors (PERI) was significantly predicted by connectivity with *rostral*-LFC regions [*r*(130) = .245, *p* = .005], confirming a link to proactive control, whereas *caudal*-LFC connectivity did not relate to PERI (see Table 1 and Figure S3). A Fisher’s r-to-z transformation demonstrated that the correlation coefficient associating *rostral*-LFC and PERI was significantly different from the coefficient for *caudal*-LFC and PERI, (Z = 2.04, *p* = .041), suggesting PERI was exclusively predicted by *rostral*-LFC connectivity. In contrast, the more simple and automatic form of post-error behavior, PES, was significantly predicted by *caudal*-LFC connectivity [*r*(130) = .244, *p* = .005], whereas *rostral*-LFC connectivity did not significantly relate to PES (see Table 1 and Figure S3), albeit the correlation coefficients for PES and caudal/rostral LFC did not significantly differ from each other (Z = 1.09 *p* = .278). Table 1 reports correlation test statistics; all significant correlations survived an FDR correction for multiple comparisons and qualitatively similar results were obtained when not collapsing across hemisphere.

**Table 1.** Post-error connectivity and post-error behavior. Pearson product-moment correlation test statistics, their significance (superscript), and corresponding sample size (italics), for a series of correlation tests between rostral/caudal lateral-frontal cortex (LFC) connectivity and either post-error reduction in interference (PERI) or post-error slowing (PES). Connectivity measures reflect MFC-seeded inter-channel phase synchrony within the post-response theta factor for the error minus correct contrast.

|  | **PERI** | **PES** |
| --- | --- | --- |
| **Rostral-LFC** | .245* | .159 |
|  | *130* | *129* |
| **Caudal-LFC** | .085 | .244* |
|  | *130* | *130* |

\* Denotes $p < .05$ after FDR correction for multiple comparisons; Post-error reduction in interference (PERI); Post-error slowing (PES); lateral-frontal cortex (LFC)

#### Conflict monitoring and pre-response reactive control

A paired-samples t-test investigating the pre-response theta factor revealed an increase in MFC total power [*t*(1,128) = 3.34 *p* = .001] prior to incongruent-correct compared to congruent-correct responses, consistent with a role in conflict monitoring. The ANOVA model investigating connectivity within this same pre-response theta factor yielded a main effect of caudality [*F*(2,236) = 54.08, *p* < .001], as well as a caudality-by-congruency interaction [*F*(2,236) = 10.47, *p* < .001]. Follow-up comparisons, collapsing across hemisphere, demonstrated significantly stronger connectivity between MFC and caudal-LFC regions, in comparison to occipital [*t*(1,116) = 2.19, *p* = .03], whereas connectivity magnitude for caudal-LFC did not significantly differ from rostral-LFC [*t*(1,116) = 1.41, *p* = .161]; connectivity strength did not differ for rostro-LFC and occipital [*t*(1,116) = 1.05, *p* = .298]. An interaction between hemisphere and congruency was also identified [*F*(1,118) = 4.14, *p* = .044], such that stronger MFC connectivity with the right hemisphere overall was observed prior to incongruent-correct responses. Collectively, this pattern of results is consistent with MFC-related conflict monitoring (theta power) recruiting LFC-related reactive control (MFC-LFC connectivity) to resolve conflict, with an emphasis on the role of more caudal-LFC regions and a right-lateralized network. Figure 4 depicts the congruent/incongruent pre-response theta power and connectivity results.

**Figure 4.**
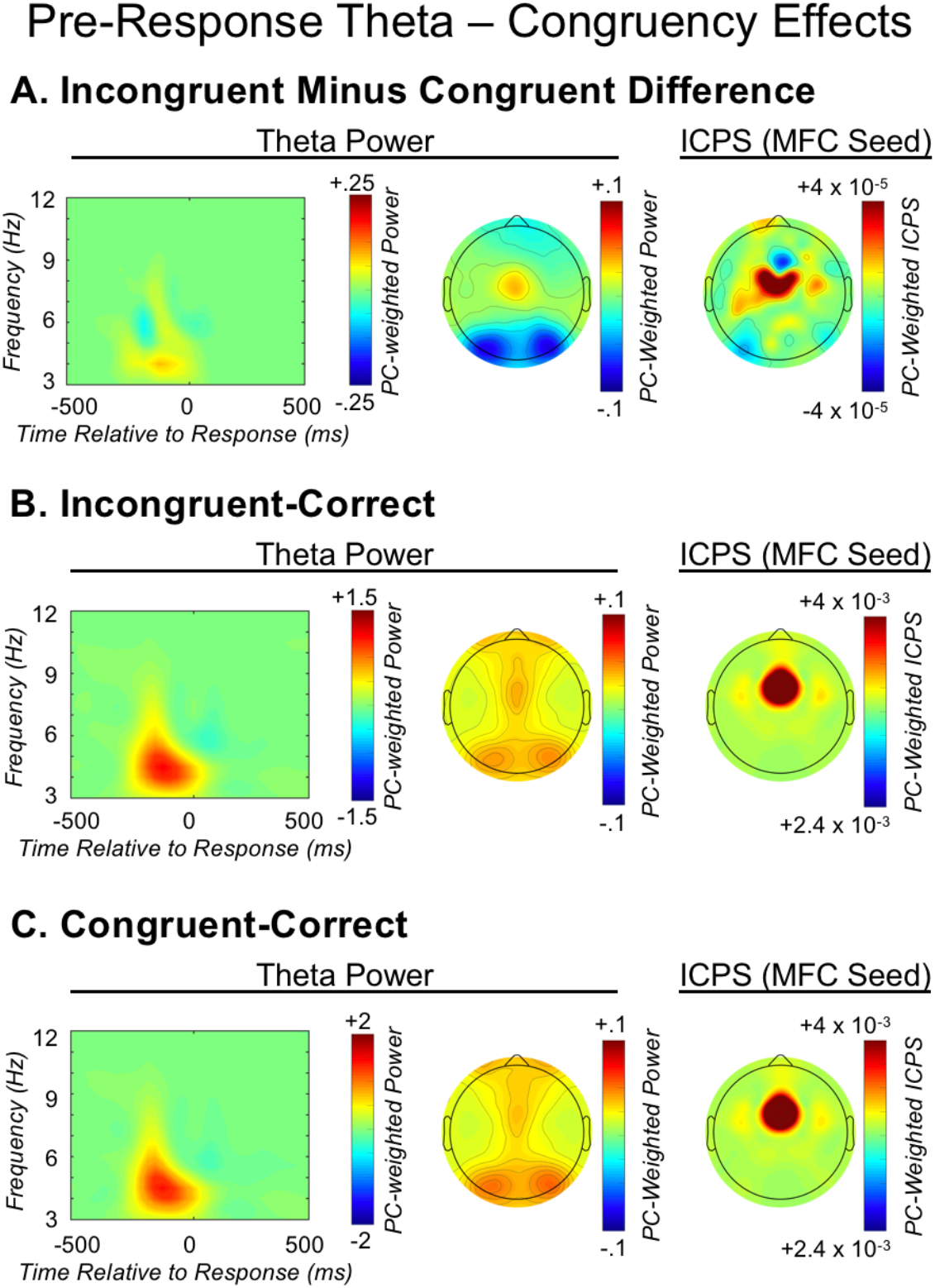
Pre-response theta dynamics. From left to right, each row depicts: the medial-frontal cortex (MFC) total power time-frequency distribution weighted by the pre-response theta factor; the corresponding topographic plot; MFC-seeded inter-channel phase synchrony (ICPS) within the pre-response theta factor. The three rows present: A) the difference between incongruent-correct and congruent-correct activity; B) incongruent-correct activity; congruent-correct activity.

To provide further evidence that pre-response MFC-LFC connectivity reflects reactive control, we tested whether such connectivity was *weaker* on *error*-incongruent trials (in comparison to *correct*-incongruent trials), suggesting a failure of reactive control leading to errors. This ANOVA model revealed a main effect of accuracy [*F*(1,120) = 27.33, *p* < .001], caudality [*F*(2,240) = 52.37, *p* < .001], and a caudality-by-accuracy interaction [*F*(2,240) = 9.02, *p* < .001]. Follow-up comparisons, collapsing across hemisphere, demonstrated that the MFC region exhibited a significant *decrease* in connectivity prior to error responses for rostral-LFC [*t*(1,120) = -4.72, *p* < .001] and caudal-LFC regions [*t*(1,120) = -4.87, *p* < .001], but did not differ for occipital regions [*t*(1,120) = -.56, *p* = .58]. These data provide further support for pre-response MFC-LFC connectivity as an index of reactive control, with *reduced* reactive control recruitment associating with error responses. Figure 5 depicts the error/correct pre-response theta connectivity results. Also note the inverse pattern of control recruitment before and after response execution: *reduced* reactive control before the response leads to errors and post-response *increases* in proactive control (for the next trial), whereas *increased* reactive control before the response leads to correct responses and reduced post-response proactive control (for the next trial). This complete cascade of cognitive control processing is depicted in Figure 6.

**Figure 5.**
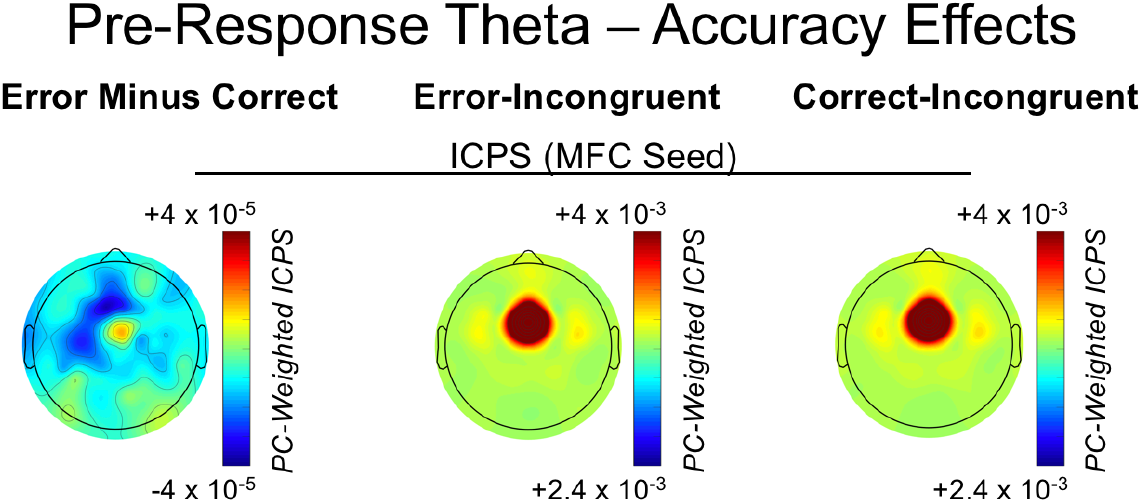
Pre-response theta dynamics as a function of response accuracy. From left to right, plots reflect medial-frontal cortex (MFC) seeded inter-channel phase synchrony (ICPS) within the pre-response theta factor for: the difference between error-incongruent and correct-incongruent activity; error-incongruent activity; correct-incongruent activity.

**Figure 6.**
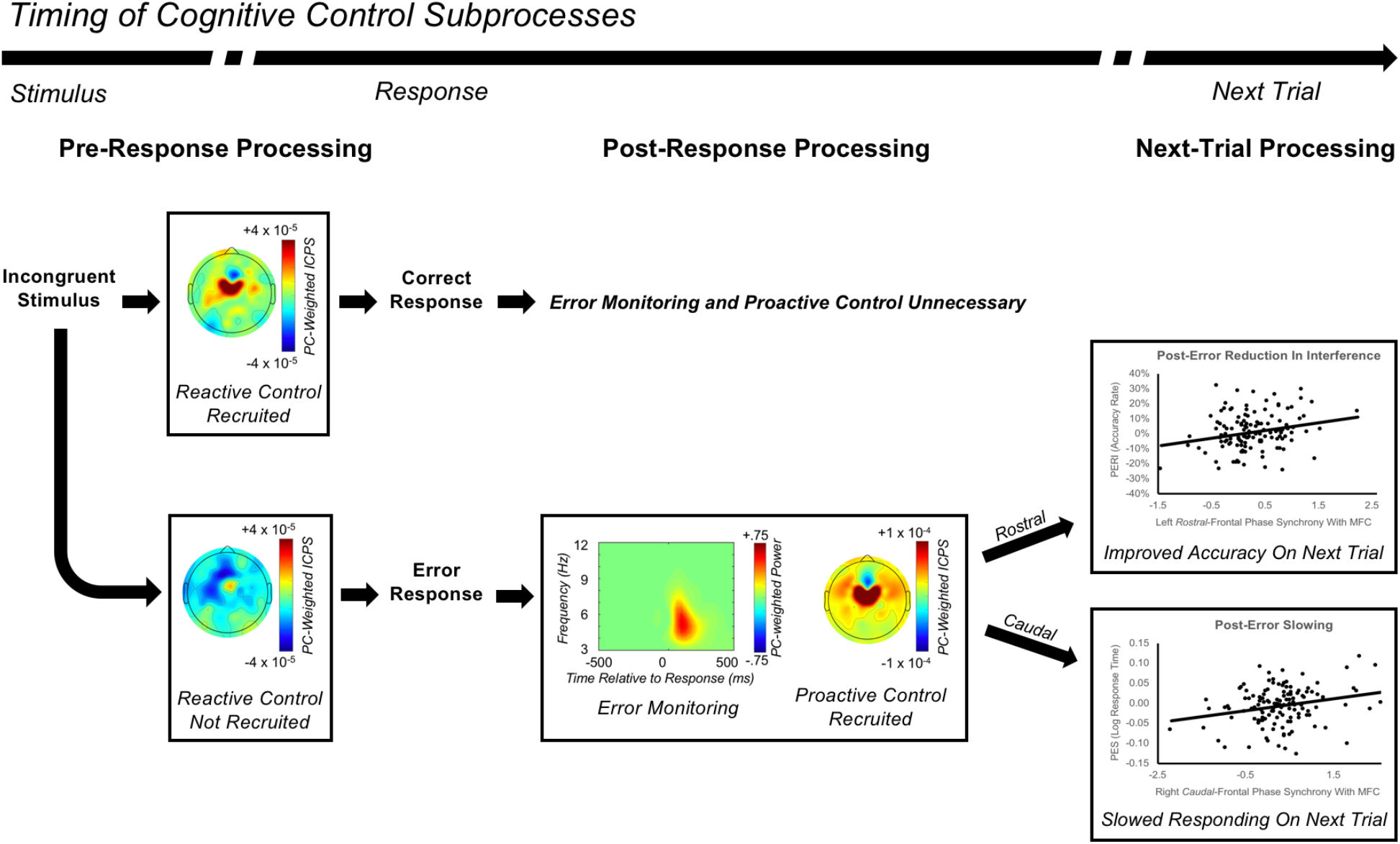
Cascade of processes involved in cognitive control. Progressing from left to right, the image depicts the relative timing of a cascade of cognitive control subprocesses. Following the presentation of a stimulus requiring control (e.g. an incongruent stimulus), if reactive control is properly recruited prior to the response then a correct response will be made, and no post-response error monitoring or proactive control (for the next trial) will be observed (top panel). In contrast, if reactive control is not properly recruited prior to the response then an error will be made, engaging error monitoring and proactive control processes (bottom panel). In turn, proactive control will influence behavior on the following trial, with more rostral-lateral-frontal cortex (LFC) regions driving post-error reduction in interference (PERI: top scatterplot), and more caudal-LFC regions driving post-error slowing (PES; bottom scatterplot).

### Effects of Social Motivation

#### Subjective reports of motivation

Consistent with our previously published study (Barker et al., 2018), participants reported significantly higher levels of effort during the social, compared to non-social, condition [*t*(1,143) = 7.353, *p* < .001]. Moreover, increased effort within the social condition was commonly attributed to social factors in the free-response explanations of effort; for example: “I tried hard because other people were giving me feedback.” Collectively, the subjective reports of motivation confirm that the social manipulation successfully increased motivation.

#### Behavior

For all analyses (behavioral and EEG), outliers (+/-3 SD) for specific conditions were removed where appropriate; each statistical analysis was performed using as much data as possible as opposed to listwise deletion across all analyses. Social motivation improved task performance, with a reduced conflict effect for accuracy in the social (compared to nonsocial) condition [*t*(1,142) = -2.65, *p* = .009]. Improved performance did not occur at the cost of a speed-accuracy trade-off, given that conflict-effect-RT did not differ between the social and nonsocial conditions [*t*(1,142) = 1.17, *p* = .243]. Similarly, social motivation did not significantly influence post-error behavior for either PERI [*t*(1,140) = -.23, *p* = .816] or PES [*t*(1,138) = .72, *p* = .474].

#### Error detection and post-error proactive control

Social motivation yielded an increase in error monitoring, with post-response MFC theta power for errors being greater in the social condition, [*t*(1,127) = 3.1, *p* = .002]. Additionally, an ANOVA model investigating post-response error-related connectivity revealed a main effect of social context, with social motivation driving an overall increase in connectivity with the MFC region [*F*(1,115) = 4.92, *p* = .028]. A main effect of caudality was also identified, with connectivity being stronger at more rostral locations [*F*(2,230) = 12.7, *p* < .001]. Finally, a post-hoc contrast identified a significant linear caudality-by-social-context interaction, such that social motivation was associated with increasingly stronger connectivity between MFC and more anterior regions (i.e. rostral- and caudal-LFC, relative to occipital) [*F*(1,115) = 4.23, *p* = .042]. Collectively, this pattern of results demonstrates that social motivation yields increases in error monitoring and post-response proactive control. The right side of Figure 7 depicts the effects of social motivation on post-response theta power and connectivity.

**Figure 7.**
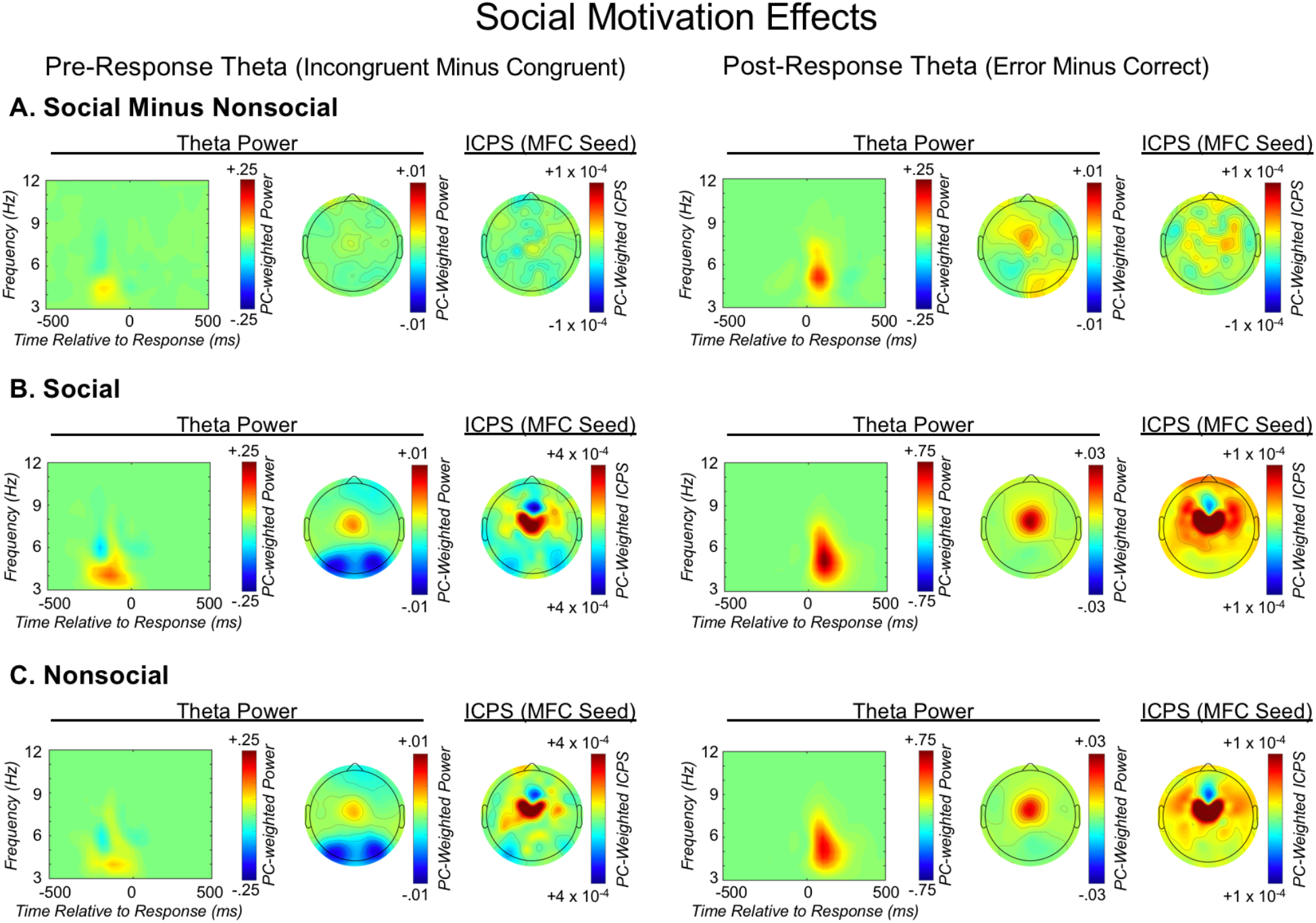
The effects of social motivation on pre- and post-response theta dynamics. From left to right, each row depicts: the medial-frontal cortex (MFC) total power time-frequency distribution weighted by the pre-response theta factor; the corresponding topographic plot; MFC-seeded inter-channel phase synchrony (ICPS) within the pre-response theta factor; the MFC total power time-frequency distribution weighted by the post-response theta factor; the corresponding topographic plot; MFC-seeded ICPS within the post-response theta factor. From top to bottom, each row depicts: A) the difference between social and nonsocial congruency-related and error-related difference scores of neural activity; B) social congruency-related and error-related difference scores of neural activity; C) nonsocial congruency-related and error-related difference scores of neural activity.

#### Conflict monitoring and pre-response reactive control

In contrast to an effect of social motivation on error monitoring, a paired samples t-test identified no effect of social motivation on conflict monitoring, as measured by pre-response and congruency-related MFC theta power [*t*(1,127) = .499, *p* = .618]. Similarly, social motivation did not influence pre-response *reactive* control, as an ANOVA model investigating congruency-related pre-response connectivity with MFC only exhibited the already described main effect of caudality [*F*(2,240) = 17.34, *p* < .001], as well as a main effect of hemisphere [*F*(1,120) = 5.46, *p* = .021] such that connectivity was increased for the right hemisphere, but this did not interact with social context. Null relations were also identified when probing pre-response connectivity as a function of accuracy, with the only significant result being the already described main effect of caudality [*F*(2,242) = 7.8, *p* = .001]. The left side of Figure 7 depicts the effects of social motivation on pre-response theta power and connectivity. Collectively these data suggest that social motivation exclusively influences post-response error monitoring and proactive control, not pre-response conflict monitoring or reactive control.

## Discussion

The current study provides a detailed characterization of cognitive control subprocess supported by theta oscillations during adolescence. At a broad level, findings for adolescent theta dynamics appeared qualitatively similar to findings in adults, with increased MFC theta power for errors and conflict monitoring, and increased MFC-LFC phase synchrony (connectivity) for proactive and reactive control recruitment. However, by leveraging Cohen’s Class RID, time-frequency PCA, and a Laplacian transform, we elucidated novel phenomena not previously observed in adolescents or adults. We dissociated pre- and post-response theta, linking greater pre-response MFC-LFC connectivity to reactive control and correct responding on the current trial. Post-response MFC-LFC connectivity exhibited the opposite pattern, being increased after errors and reflecting proactive control for the following trial. We further characterized differences in MFC connectivity with rostral/caudal LFC after errors: MFC connectivity with rostral-LFC predicted post-error reduction in interference (PERI), whereas connectivity with caudal-LFC predicted post-error slowing (PES). We leveraged this improved characterization of cognitive control—afforded by time-frequency EEG—to distinguish *how* social motivation influences adolescent cognitive control. Social motivation exclusively upregulated post-response error monitoring (MFC theta power) and proactive control (MFC-LFC connectivity), but not pre-response conflict monitoring and reactive control. Collectively, the current study details the role of theta oscillations and social motivation on adolescent cognitive control, while informing research on cognitive control more generally.

In adults, evidence suggests MFC neurons subserve performance monitoring (Ridderinkhof et al., 2004), with LFC instantiating top-down control (Kerns et al., 2004). fMRI has confirmed similar MFC and LFC roles during adolescence (Crone and Steinbeis, 2017). However, adult studies use measures of theta to index unique organizing properties of cognitive control (Cavanagh and Frank, 2014; Verguts, 2017); the current results extend such work to adolescence. Conflict and error monitoring were associated with increased MFC theta power, and control recruitment relied on MFC-LFC theta phase synchrony (ICPS; connectivity). Crucially, we demonstrated theta’s behavioral relevance, with accurate responses linked to pre-response MFC-LFC phase connectivity, and post-error behavioral changes linked to post-response MFC-LFC theta phase connectivity. These findings are consistent with prior results in adult humans, non-human primates (Tsujimoto et al., 2006) and other mammals (Narayanan et al., 2013), but also provide novel insights regarding the distinction between pre- and post-response theta and link them to the concept of reactive/proactive control. This work provides a foundation for mechanistic accounts of adolescent cognitive control and its relation to motivational factors during a critical window of human development.

The current results also extend prior work suggesting that cognitive control develops non-linearly and approaches maturity in adolescence (Casey et al., 2001; Luna et al., 2004). Children differ markedly from adults in behavior (Casey et al., 2001; Luna et al., 2004), ERP measures of error monitoring (Davies et al., 2006), and interconnectivity between medial and lateral frontal cortex (Hwang et al., 2010). However, adolescents often reach adult-levels of behavioral performance (Casey et al., 2001; Luna et al., 2004), particularly when incentivized (Padmanabhan et al., 2011), demonstrate an absence of changes in the core error-monitoring system and its relations with post-error control (Buzzell et al., 2017b), and exhibit a majority of medial-lateral frontal connections similar to those of adults (Hwang et al., 2010).

Whereas adolescent cognitive control can appear adult-like in some contexts, motivational factors may disproportionately affect adolescents (Casey et al., 2008; Luciana and Collins, 2012; Luna et al., 2015; Nelson et al., 2005; Steinberg et al., 2008). Indeed, this effect can obscure adult-like functions in some contexts (Luciana and Collins, 2012), which generates much interest and debate (Crone and Dahl, 2012; Pfeifer and Allen, 2012; Shulman et al., 2016). Here, we illustrate novel features of these motivation-control relations by parsing cognitive control into particular subprocesses. Social motivation exclusively influenced post-response error monitoring (theta power) and proactive control (MFC-LFC connectivity). Similarly, social motivation yielded task-specific accuracy improvements (reduced behavioral conflict-effect). This pattern matches effects induced by monetary incentives on proactive control in adults (Botvinick and Braver, 2015) and may explain other findings on motivational influences during adolescence (Luciana et al., 2012). The current study demonstrates heterogeneity in motivation-control relations during early adolescence to be further understood through future studies.

The current report also provides critical extensions of prior adult work regarding the neural mechanisms involved in cognitive control. fMRI studies suggest a rostro-caudal order regarding LFC control function (Badre, 2008; Badre and Wagner, 2004; Koechlin, 2003), with caudal regions associated with simpler forms of control, including motor inhibition, and rostral regions linked to more complex forms of control (Badre and Wagner, 2004; Koechlin, 2003). However, different forms of control are known to follow errors (Danielmeier and Ullsperger, 2011), and yet, rostral-caudal theories have not previously been applied to explain such findings. Here, we use MFC theta phase synchrony (connectivity) to provide the first evidence distinguishing forms of post-error behavioral control related to rostral/caudal LFC. MFC connectivity with caudal-LFC predicted PES, a relatively automatic and general change in behavior associated with motor inhibition (King et al., 2010; Notebaert et al., 2009; Wessel and Aron, 2017), whereas MFC connectivity with rostral-LFC predicted PERI, a more deliberative and task-specific form of proactive control linked to selective attention (King et al., 2010; Maier et al., 2011; Ridderinkhof, 2002). Clearly, these findings require replication in adults. Nevertheless, these finding may suggest a way to integrate understandings in prior research on distinct features of cognitive control across three levels of analysis: 1) theory-based distinctions between MFC for monitoring and LFC for control (MacDonald et al., 2000), 2) anatomy-based distinctions based on rostral-caudal LFC organization (Badre, 2008), and 3) neurophysiology-based distinctions based on theta as an organizing rhythm for cognitive control (Cavanagh and Frank, 2014).

These data also inform debates regarding whether error processing is adaptive (Danielmeier and Ullsperger, 2011; Jentzsch and Dudschig, 2009; Notebaert et al., 2009; Wessel, 2018). PES is the most widely reported post-error behavioral phenomenon and traditionally viewed as an adaptive response reflecting increased cautiousness after errors (Botvinick et al., 2001). However, this view has been challenged, as PES often does not always relate to improved accuracy or attention (Buzzell et al., 2017c; Jentzsch and Dudschig, 2009; Notebaert et al., 2009). Even when not maladaptive, PES reflects a general response to *all* post-error trials, contrasting with PERI, which is task-specific and an unequivocal correlate of adaptive behavior and proactive control (Danielmeier and Ullsperger, 2011; King et al., 2010; Maier et al., 2011; Ridderinkhof, 2002). An emerging view is that both adaptive and non-adaptive responses can be observed following errors (Danielmeier and Ullsperger, 2011; Purcell and Kiani, 2016; Wessel, 2018). The current data support this view by demonstrating how two distinct forms of post-error behavior can emerge *in parallel* after errors: simultaneous communication between a common MFC node and separate LFC subregions through communication channels supported by synchronized theta oscillations.

A combination of Cohen’s class RID and time-frequency PCA (Bernat et al., 2005) allowed isolation of explicitly pre- and post-response theta dynamics *within the same epochs*. This separation removed a confound present in fMRI studies, where pre- and post-response cognitive control likely blends together due to sluggishness of the BOLD signal. However, this confound also plagues traditional time-frequency EEG studies because the limited resolution of Morlet wavelets (Bernat et al., 2005) that impairs separation of pre- and post-response theta. The current study details separation of pre- and post-response theta dynamics and their differential relations with task behavior, demonstrating the functional importance of isolating these processes. Whereas pre-response connectivity was related to correct responses on the current trial (reactive control), post-response connectivity related to improved performance on the following—post-error—trial (proactive control). Critically, we also demonstrate that pre- and post-response control are inversely related. Pre-response failures of reactive control (reduced MFC-LFC connectivity) were associated with error responses, followed by post-response connectivity increases reflecting transient proactive control to adapt future behavior. Conversely, successful pre-response reactive control yielded correct responses, followed by reduced post-response connectivity. The Cascade of Control Model (Banich, 2009) proposes similar inverse relations before/after stimulus presentation, with pre-stimulus control reducing the need for reactive control after stimulus presentation. However, the current results present the first evidence for similar inverse relations before/after *response execution*. Thus, separation of pre- and post-response theta allowed for galvanizing the notion of inverse control relations into a more general principle of the cognitive control system. However, similar observations in adults are needed.

Collectively, the current report provides a detailed account of theta oscillations in adolescent cognitive control, elucidates the nuanced effects of social motivation, and identifies several novel mechanisms of cognitive control more broadly. Linking adolescent cognitive control to theta dynamics opens the door to theoretical integration across developmental stages and even species, complementing existing fMRI/ERP studies of adolescent cognitive control. Identifying dissociations in *how* social motivation influences control can inform future investigations into motivation-control relations during this critical period of human development. Finally, the methodology employed here can be leveraged by future studies to more broadly characterize typical and atypical cognitive control dynamics across age.

## Acknowledgements

This research was supported by grants from the National Institute of Mental Health (U01MH093349 and U01MH093349-S to NAF; P01HD064653 supporting EMB), the National Science Foundation (DGE1322106 to SVT), the Eunice Kennedy Shriver National Institute of Child Health and Human Development (5T32HD007542 supporting TVB), and the NIMH-Intramural Research Program (ZIAMH-002782 supporting DSP). The authors declare no competing financial interests.

## Author Contributions

Conceptualization, G.A.B., H.A.H., L.C.B., N.A.F., S.V.T.R. and T.V.B.; Methodology, L.C.B., S.V.T.R. and T.V.B.; Investigation, L.C.B., M.E.B, S.V.T.R. and T.V.B.; Data Curation, G.A.B., S.M. and T.V.B.; Software, G.A.B., E.M.B. and S.M.; Formal Analysis, G.A.B., E.M.B. and M.E.B.; Visualization, G.A.B.; Writing – Original Draft, G.A.B.; Writing – Review & Editing, G.A.B., D.S.P., E.M.B., H.A.H., L.C.B., M.E.B., N.A.F., S.M., S.V.T.R. and T.V.B.; Resources, N.A.F.; Project Administration, N.A.F.; Funding; Acquisition, H.A.H., N.A.F.; Supervision, H.A.H., D.S.P., N.A.F.

## Declaration of Interests

The authors declare no competing interests

---

**Figure S1.**
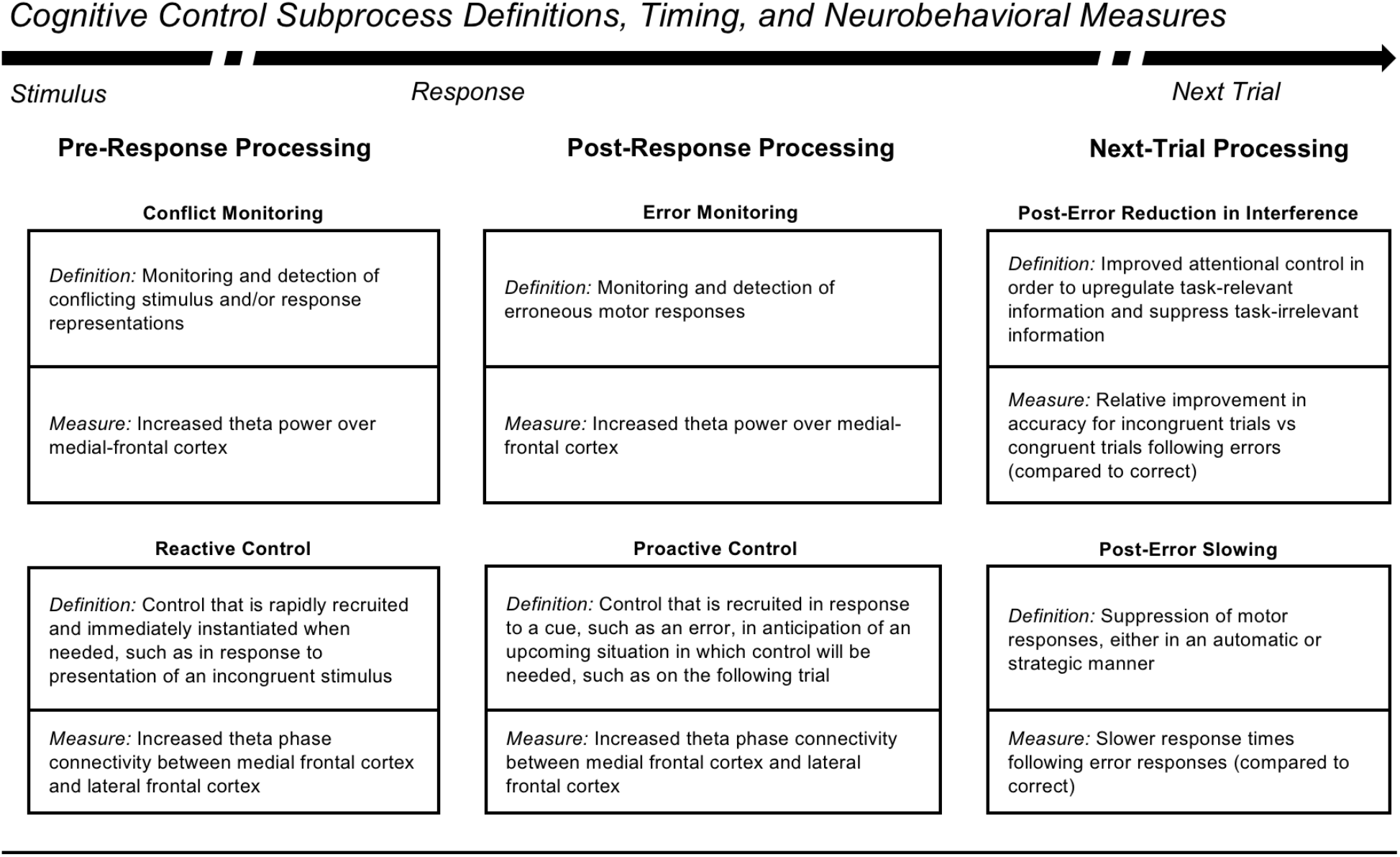
Description of cognitive control subprocesses and neurobehavioral measures. The arrow depicts the flow of time for a single trial on a task requiring cognitive control (e.g. a flanker task). Time begins with stimulus presentation and pre-response processing, followed by response commission and post-response processing, ending with presentation of a subsequent trial and associated neurobehavioral processing. Each box defines a particular cognitive control subprocess and a neural or behavioral measure that can be used to index the subprocess. Note that the use of proactive control here is distinct from the more common study of “tonic proactive control” that occurs at the block level. Instead, our use of proactive control is in line with the notion of “transient proactive control” that can follow an error and prepare control for the subsequent trial in a proactive manner. See Table S1 for definitions.

**Table S1.**
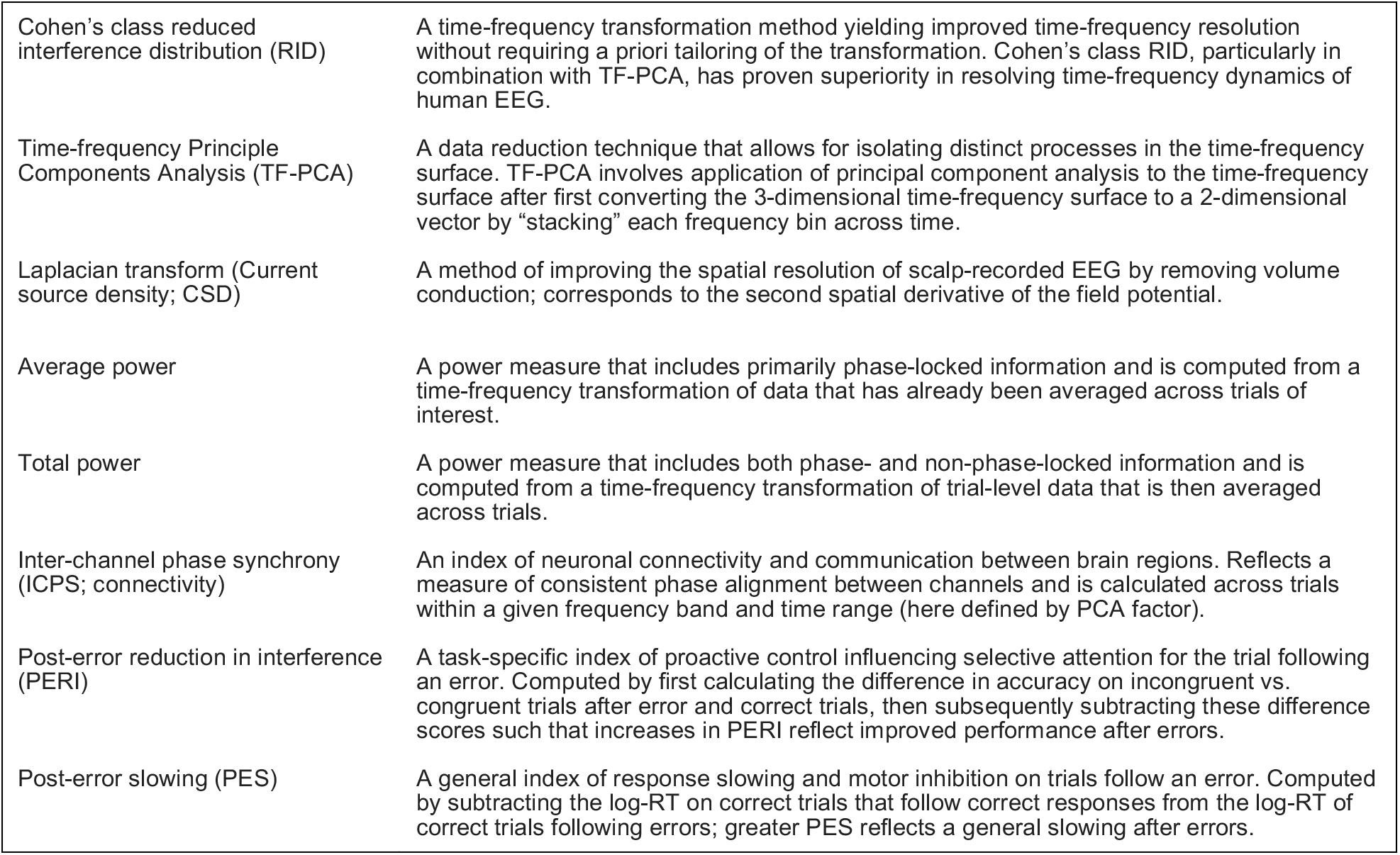
Important terms and definitions.

|  |  |
| --- | --- |
| Cohen's class reduced interference distribution (RID) | A time-frequency transformation method yielding improved time-frequency resolution without requiring a priori tailoring of the transformation. Cohen's class RID, particularly in combination with TF-PCA, has proven superiority in resolving time-frequency dynamics of human EEG. |
| Time-frequency Principle Components Analysis (TF-PCA) | A data reduction technique that allows for isolating distinct processes in the time-frequency surface. TF-PCA involves application of principal component analysis to the time-frequency surface after first converting the 3-dimensional time-frequency surface to a 2-dimensional vector by "stacking" each frequency bin across time. |
| Laplacian transform (Current source density; CSD) | A method of improving the spatial resolution of scalp-recorded EEG by removing volume conduction; corresponds to the second spatial derivative of the field potential. |
| Average power | A power measure that includes primarily phase-locked information and is computed from a time-frequency transformation of data that has already been averaged across trials of interest. |
| Total power | A power measure that includes both phase- and non-phase-locked information and is computed from a time-frequency transformation of trial-level data that is then averaged across trials. |
| Inter-channel phase synchrony (ICPS; connectivity) | An index of neuronal connectivity and communication between brain regions. Reflects a measure of consistent phase alignment between channels and is calculated across trials within a given frequency band and time range (here defined by PCA factor). |
| Post-error reduction in interference (PERI) | A task-specific index of proactive control influencing selective attention for the trial following an error. Computed by first calculating the difference in accuracy on incongruent vs. congruent trials after error and correct trials, then subsequently subtracting these difference scores such that increases in PERI reflect improved performance after errors. |
| Post-error slowing (PES) | A general index of response slowing and motor inhibition on trials follow an error. Computed by subtracting the log-RT on correct trials that follow correct responses from the log-RT of correct trials following errors; greater PES reflects a general slowing after errors. |

**Figure S2.**
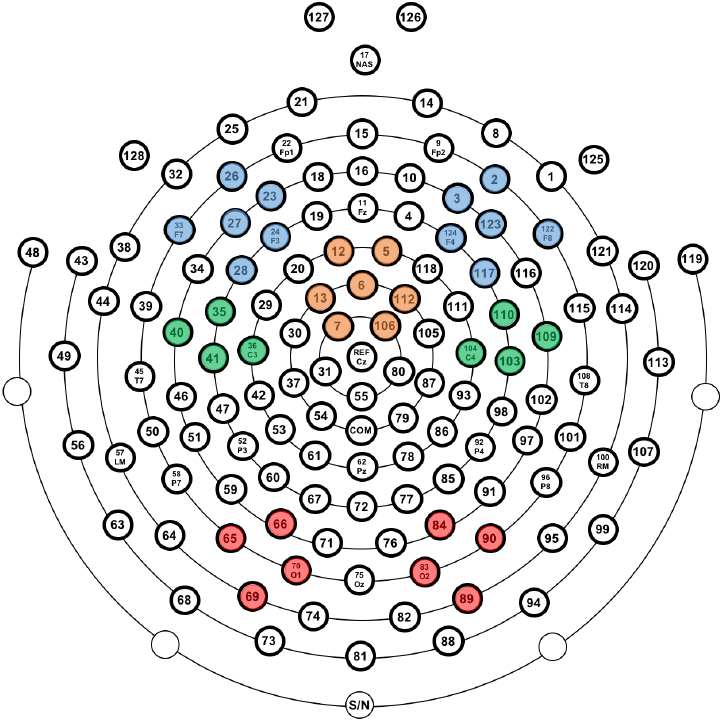
Electrode clusters employed in all EEG analyses. Medial-frontal cortex (orange); left and right rostral-lateral-frontal cortex (blue); right and left caudal-lateral-frontal cortex (green); right and left occipital cortex (red).

**Figure S3.**
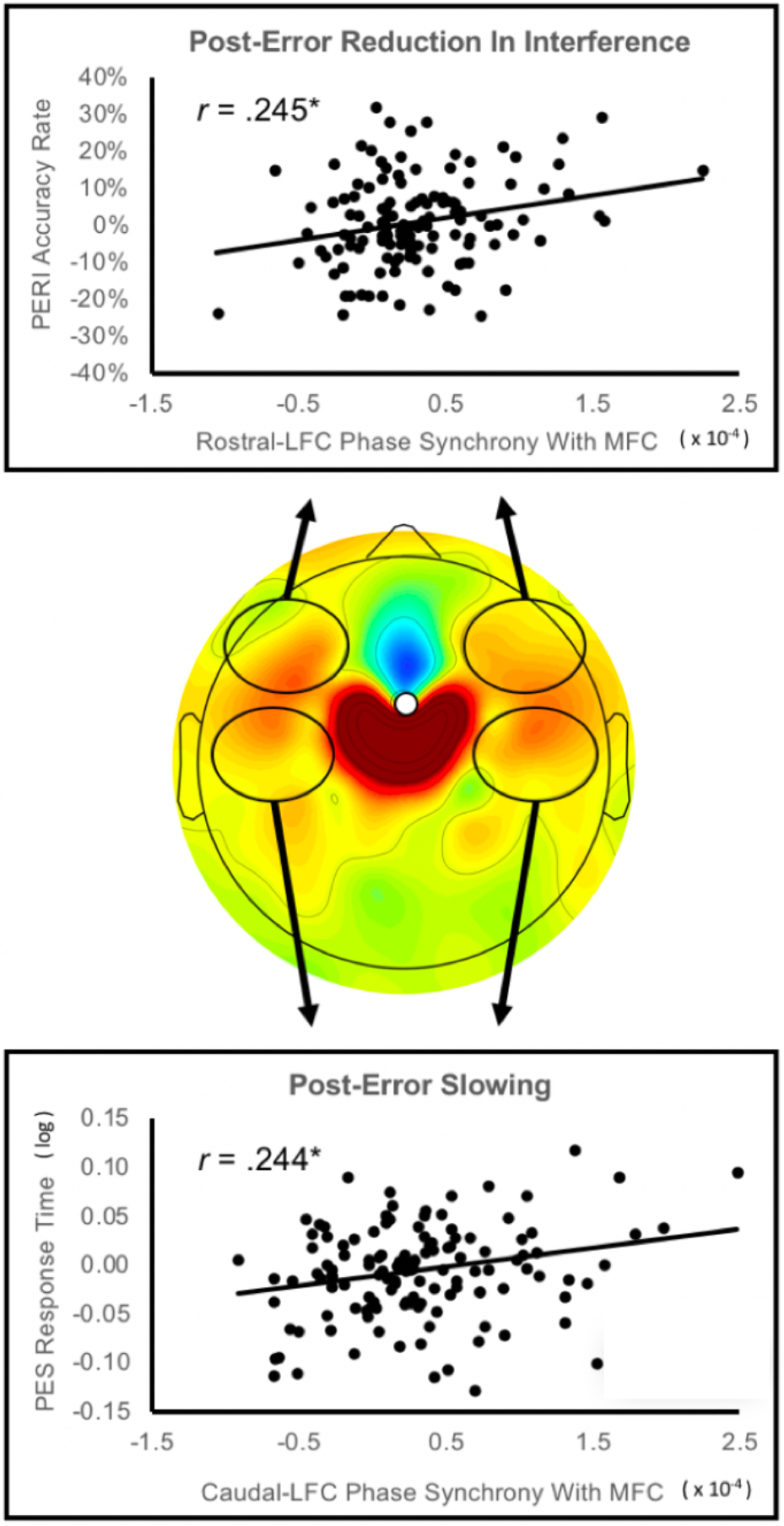
Relations between post-error MFC-LFC connectivity and next-trial behavior. The central plot depicts the increase in medial-frontal cortex (MFC) to lateral-frontal cortex (LFC) connectivity within the post-response theta factor (MFC seed; error minus correct difference); black ellipses indicate the location of electrode clusters used to quantify MFC connectivity with rostral/caudal LFC. The top scatterplot depicts relations between bilateral *rostral*-LFC connectivity and post-error reduction in interference (PERI); the bottom scatterplot depicts relations between bilateral *caudal*-LFC connectivity and post-error slowing (PES).

